# Acrylamide Inhibits Vaccinia Virus Through Vimentin-independent Anti-Viral Granule Formation

**DOI:** 10.1101/2020.07.30.228858

**Authors:** Jennifer J Wood, Ian J White, Jason Mercer

## Abstract

The replication and assembly of vaccinia virus (VACV), the prototypic poxvirus, occurs exclusively in the cytoplasm of host cells. While the role of cellular cytoskeletal components in these processes remains poorly understood, vimentin - a type III intermediate filament - has been shown to associate with viral replication sites and to be incorporated into mature VACV virions. Here we employed chemical and genetic approaches to further investigate the role of vimentin during the VACV lifecycle. The collapse of vimentin filaments, using acrylamide, was found to inhibit VACV infection at the level of genome replication, intermediate- and late- gene expression. However, we found that CRISPR-mediated knockout of vimentin did not impact VACV replication. Combining these tools, we demonstrate that acrylamide treatment results in the formation of antiviral granules (AVGs) known to mediate translational inhibition of many viruses. We conclude that vimentin is dispensable for poxvirus replication and assembly and that acrylamide, as a potent inducer of AVGs during VACV infection, serves to bolster cell’s antiviral response to poxvirus infection.

**Summary Statement:** Acrylamide inhibits poxvirus replication by inducing anti-viral granules and blocking translation. This inhibition is independent of the effect of acrylamide on vimentin filaments which were found to be dispensable for viral replication and assembly.

## Introduction

Poxviruses are a family of large dsDNA viruses that replicate exclusively in the cytoplasm of host cells (Moss 2013). Being the most complex mammalian viruses, they engage with a large repertoire of host proteins and processes to ensure successful infection and spread (Mercer et al. 2012). As the prototype member of this family, vaccinia virus (VACV), has become a useful to tool for investigation of basic virus-host interactions and for the development of anti-poxviral strategies (Bidgood 2019).

It is well accepted that VACV modulates the two main components of the host cytoskeleton (actin and microtubules) throughout its lifecycle. Dramatic reorganisation of host cell actin occurs during virus entry (Mercer and Helenius 2008, Mercer et al. 2010) and egress (Smith and Law 2004, Cudmore et al. 1995), and microtubules are subjugated to coalesce transcription and replication sites (Mallardo, Schleich and Locker 2001), to transport nascent virions from assembly to wrapping compartments and subsequently to the cell surface (Ward and Moss 2001, Rietdorf et al. 2001).

It has also been reported that VACV infection results in reorganisation of vimentin, a type III intermediate filament involved in a range of cellular functions including; cell migration, proliferation, signal transduction and the organisation of cytosolic organelles (Danielsson et al. 2018, Chang and Goldman 2004, Ivaska et al. 2007, Minin and Moldaver 2008, Lowery et al. 2015, Styers et al. 2004). For VACV, vimentin filaments were shown to surround and concentrate within viral replication sites (Risco et al. 2002), leading to the hypothesis that it plays a role in their formation and in the assembly of early virion intermediates (Risco et al. 2002). In support of this hypothesis, vimentin was identified in proteomics analyses of purified mature VACV virions, suggesting that this intermediate filament was associated with, or packaged into, viral particles during assembly (Resch et al. 2007, Chung et al. 2006).

In the absence of small compounds that specifically depolymerise vimentin, acrylamide - which disrupts vimentin polymerisation without impacting other cytoskeletal elements (Eckert 1986, Durham, Pena and Carpenter 1983) - is often employed to study the interplay between viruses and vimentin. In addition to its impact on vimentin, acrylamide treatment has been shown to activate several oxidative and ER stress pathways (Komoike and Matsuoka 2016). Exposure to acrylamide results in the production of reactive oxygen species and mis-regulation of cellular redox status (Kim et al. 2015, Jiang et al. 2007) which ultimately triggers eIF2α signalling and a shift from global to stress specific protein synthesis (Komoike and Matsuoka 2016, Komoike and Matsuoka 2019, Jackson, Hellen and Pestova 2010). Using acrylamide as a tool, vimentin has been reported to play diverse roles in the lifecycles of several viruses including: Chikungunya virus (Issac, Tan and Chu 2014), HIV (Wang et al. 2016), HCMV (Miller and Hertel 2009), MVM Parvovirus (Fay and Pante 2013), Junin (Cordo and Candurra 2003) and bluetongue virus (Bhattacharya, Noad and Roy 2007).

Here we set out to investigate the role of vimentin in VACV replication and assembly. In line with previous reports (Risco et al. 2002, Resch et al. 2007), we observed that during VACV infection vimentin associates with replication sites and is packaged into newly assembled virions. We demonstrate that treatment of infected cells with acrylamide, which correlates with the collapse of vimentin filaments, dramatically impacts VACV production through inhibition of genome replication and subsequent gene expression. Surprisingly, upon generating a vimentin-null cell line we found that infection was still sensitive to acrylamide and that viral replication was not altered by the loss of vimentin. Consistent with a vimentin-independent effect, we show that acrylamide treatment leads to the formation of anti-viral granules, which appear to block post-replicative gene expression and the production of new virus particles.

## Results

### Vimentin is peripheral to VACV replication sites and within mature virions

The intermediate filament vimentin has been observed to be associated with VACV replication sites by electron and fluorescence microscopy (Risco et al. 2002, Resch et al. 2007). To verify the rearrangement and relocalisation of vimentin to sites of VACV replication, HeLa cells were infected with VACV and at 6 hours post infection (hpi) cells were stained with Hoechst to visualize viral replication sites and immunolabelled for vimentin. In uninfected control cells vimentin appeared as a tubular network localised around the nucleus of the cell (Fig 1A; top). In VACV infected cells, as previously reported, we found vimentin in close association with the periphery of multiple viral replication sites present in the cytoplasm (Fig 1A; bottom see ROI).

**Figure 1.**
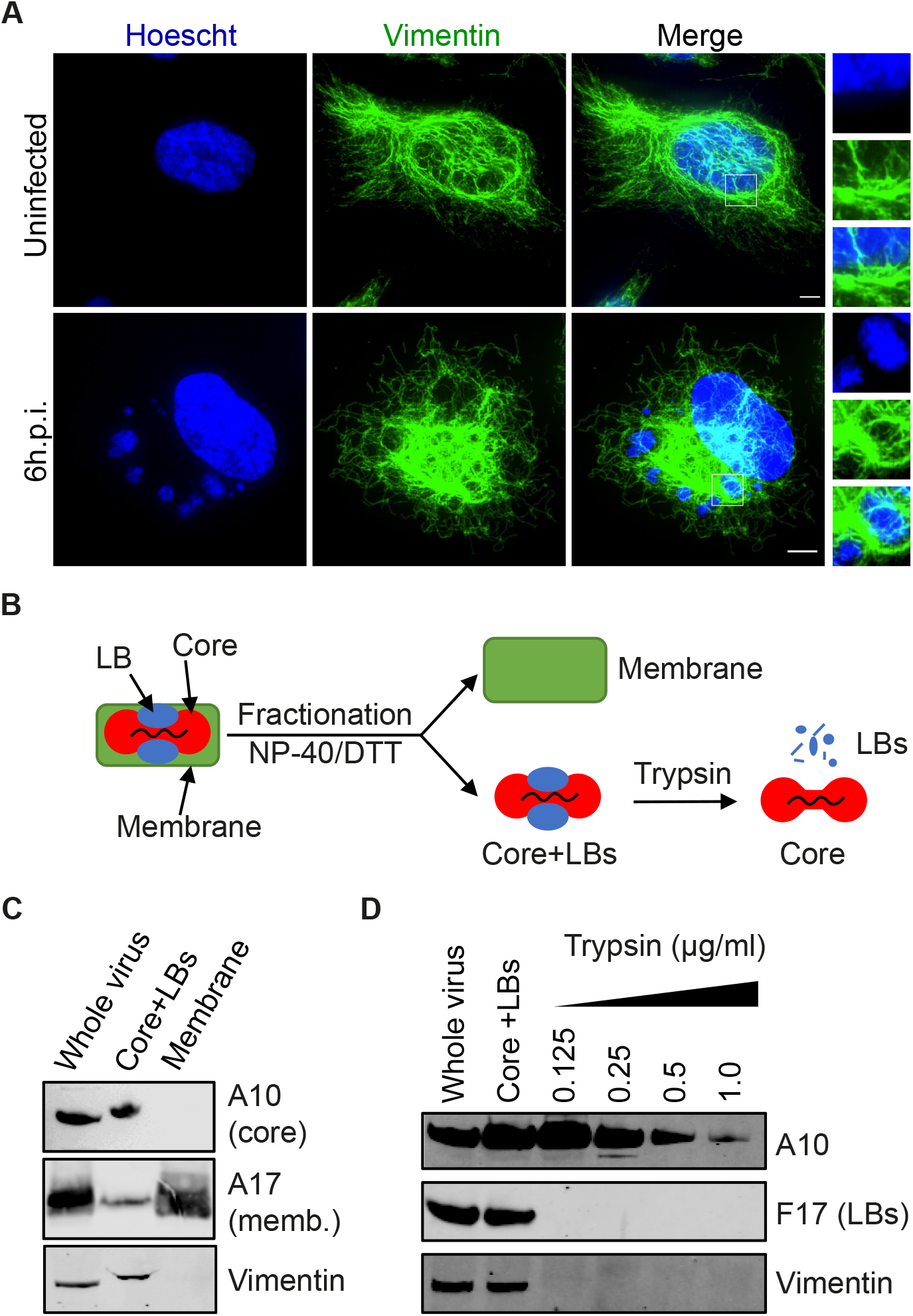
Vimentin is associated with VACV replication. (A) HeLa cells were infected with WR VACV at MOI 10 for 6 hours then fixed and stained with anti-vimentin (green). DNA visualised using Hoechst. Scale bars = 5μm. (B) Illustration of VACV fractionation protocol. Intact purified virions were subjected to treatment with detergent and reducing agents to separate viral membranes and core+LBs. To remove LBs, Core+LBs fractions were treated with trypsin (details in M and Ms). (C) Representative western blot of purified WT WR VACV virions, VACV core+LBs and membrane fractions as prepared per (B). Fractions were immunoblotted for A10 (core protein), A17 (membrane protein) and vimentin. (D) Representative western blot of purified WT WR VACV virions fractionated, and LBs removed as per (B) using increasing concentrations of trypsin. Samples were immunoblotted for A10 (core), F17 (LB) and vimentin.

Next, we sought to confirm the presence of vimentin within purified virions. While vimentin has been observed within assembling cytoplasmic virions by immuno-electron microscopy (EM) (Risco et al. 2002), its presence within mature virions (MVs) as determined by mass spectrometry is debated (Resch et al. 2007, Chung et al. 2006). To confirm and extend these findings, purified wild type VACV MVs were subjected to fractionation into core+LB and membrane samples as previously described (Mercer and Traktman 2003). Whole virions, core-LB and membrane fractions were then subjected to immunoblot analysis directed against the core protein A10, the membrane protein A17 and vimentin (Fig. 1B). Separation of A10 into core+LB and A17 into membrane samples indicated that fractionation was successful. Vimentin was present in whole virus and, upon fractionation, exclusively found in the core+LB sample (Fig. 1B; bottom panel). These results confirm the presence of vimentin in VACV virions and demonstrate its association with VACV cores or LBs.

To further refine its intra-virion localisation, we subjected the core+LB fraction to increasing trypsin treatment to digest the LBs away from the viral cores; a procedure adapted from Ishihashi and Oie (Ichihashi, Oie and Tsuruhara 1984). As expected, at low trypsin concentrations the core protein A10 remained intact, while the LB protein F17 (Schmidt et al. 2013) was completely degraded (Fig. 1C). Immunoblot analysis of vimentin mirrored that of F17, suggesting that vimentin is accessible and perhaps associated with LBs (Fig. 1C; bottom panel)

### Acrylamide inhibits VACV infection at intermediate and late transcription

Based on its localisation during infection, Risco *et al.* proposed that vimentin plays a role in the formation for VACV replication sites and virions (Risco et al. 2002). Having confirmed these localisation studies, we wanted to test the importance of an intact vimentin network during VACV infection. With its ability to collapses vimentin filaments well documented (Durham et al. 1983, Miller and Hertel 2009), we proceeded to evaluate the impact of acrylamide treatment on VACV infection. Cells were infected in the presence of acrylamide and the 24 h viral yield determined. Strikingly, the viral yield in the presence of acrylamide was reduced by greater than 3-logs (Fig. 2A). DMSO and Nocodazole, which destabilizes microtubules, were used as controls. Treatment with either did not result in significant reduction in virus production consistent with previous reports (Ploubidou et al. 2000, Rizopoulos et al. 2015). Given the dramatic reduction in virus yield seen in the presence of acrylamide, we asked which stage of the VACV life cycle was inhibited.

**Figure 2.**
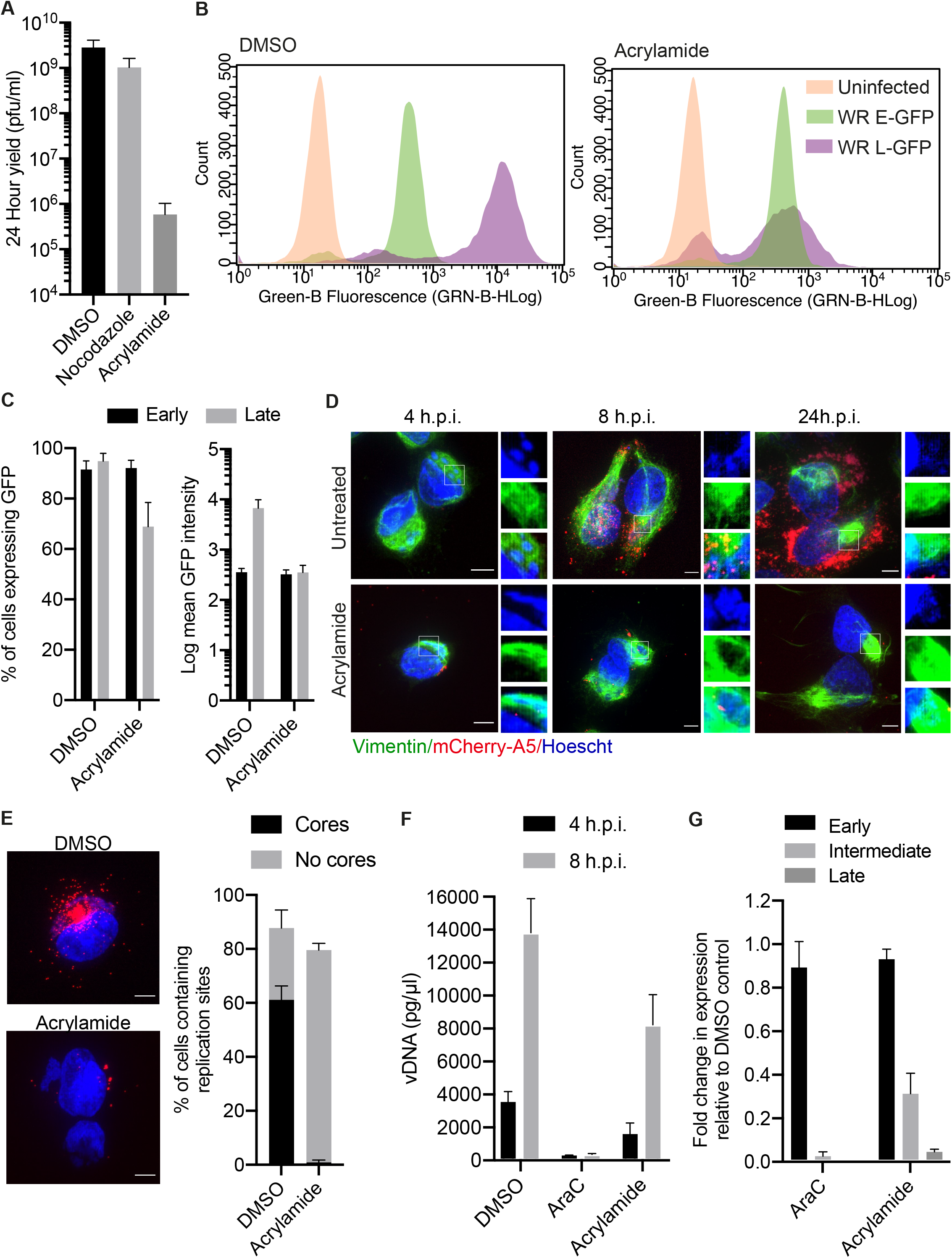
Acrylamide inhibits VACV infection. (A) HeLa cells treated with either DMSO, nocodazole or acrylamide were infected with WT WR VACV at MOI 1 and viral yield determined at 24 hpi on BSC40 cells. (B) HeLa cells treated with DMSO (left) or acrylamide (right) were infected with WR E-EGFP or L-EGFP VACV at MOI 4 and analysed by flow cytometry at 6 (early) or 8 (late) hpi. Representative flow traces of cell count vs. fluorescence intensity for uninfected (orange) WR E-EGFP (green) and WR L-EGFP (purple) cell populations. (C) Mean percentage of EGFP expressing cells and Log EGFP intensity of three independent experiments from (B). (D) HeLa cells infected with WR mCh-A5 (red) at MOI 10 in the presence of DMSO or acrylamide were fixed at 4, 8 and 24 hpi. Cells were immunostained for vimentin (green), DNA visualised using Hoechst (blue). Scale bars = 5μm (E) HeLa cells were infected with WR mCh-A5 (red) at MOI 10 in the presence of DMSO or acrylamide. At 8 hpi cells were stained for DNA using Hoechst (blue). Scale bars = 5μm. The percentage of cells containing replication sites with and without new virions was quantified (n>75). (F) To quantify VACV genome replication, HeLa cells were infected with WT WR at MOI 10 in the presence of DMSO, AraC or acrylamide. Genomic DNA (at 4 and 8 hpi) was extracted and quantified by qPCR. (G) HeLa cells were infected as in (F) and samples harvested at 2, 4 or 6 hpi. To quantify vRNA levels RTqPCR was performed using early, intermediate or late gene specific primers. GAPDH was used to normalise expression across all samples and fold change in vRNA calculated using threshold cycles. All experiments were performed in triplicate with graphs representing the mean + 1 SD.

VACV gene expression occurs in 3 temporal stages with early genes being expressed before genome replication within the virus core, and intermediate as well as late genes after genome replication (Moss 2013). To determine if the acrylamide block was pre- or post-genome replication we infected HeLa cells with recombinant viruses that express EGFP under the control of early (WR E-EGFP) or late (WR L-EGFP) viral promoters and analysed them for EGFP expression by flow cytometry (Yakimovich et al. 2017). When plotted as a histogram (cell number vs. fluorescence intensity), no defect in either the number of infected cells or intensity of EGFP expression was seen with WR E-EGFP virus (green)in the presence of acrylamide (Fig. 2B and 2C). However, a dramatic reduction in the fluorescence intensity of cells infected with WR L-EGFP (purple) was observed in the presence of acrylamide (Fig. 2B). Quantification showed that the number of infected cells was reduced by 26%, and the fluorescence intensity of infected cells reduced by 1.3 logs (Fig. 2B and 2C). That reduced levels of late gene expression were observed indicated that acrylamide blocks the VACV lifecycle after early gene expression.

To investigate post replicative stages of VACV infection HeLa cells were infected with a fluorescent recombinant virus that expresses and packages a mCherry tagged version of the VACV core protein A5 (WR mCh-A5) (Schmidt et al. 2011). Cells were left untreated or were treated with acrylamide and infection allowed to proceed for 4, 8 or 24 h prior to fixation. Nuclei and viral replication sites were visualized using Hoechst, vimentin via immunostaining and virions through the incorporation of mCh-A5. In untreated cells, at 4 hpi cytoplasmic replication sites surrounded by vimentin were seen (Fig. 2D; top). By 8 hpi, nascent virions had formed, and the vimentin appeared more loosely associated with replication sites, a phenotype that was more pronounced at 24 hpi. Conversely, infected cells treated with acrylamide displayed a total collapse of vimentin filaments around viral replication sites at all time points (Fig. 2A; bottom). Notably, in the presence of acrylamide replication sites appeared qualitatively smaller than in control cells and no new virions were formed. Quantification of the number of cells containing replication sites and VACV cores at 8 hpi showed that in the presence of acrylamide replication sites could form, but no virions were assembled (Fig. 2E).

Having observed smaller replication sites in the presence of acrylamide we next quantified its effect on viral genome replication. HeLa cells were infected with WT VACV, in the presence or absence of acrylamide, and genomic DNA extracted at 4 and 8 hpi. The amount of VACV DNA was then quantified using qPCR as previously described (Huttunen and Mercer 2019). Cytosine arabinoside (AraC), a known inhibitor of VACV DNA replication was included as a control (Balzarini and Declercq 1989, Sidwell et al. 1968). As expected, there was a ~4-fold increase in VACV DNA between 4 and 8 hpi in control cells, and only negligible amounts of vDNA in AraC treated samples (Fig. 2F). We found that acrylamide did not completely inhibit VACV DNA accumulation but reduced the levels of VACV DNA, relative to DMSO controls, at both timepoints (Fig. 2F). These results are consistent with our immunofluorescence images and suggest that acrylamide partially blocks or slows VACV DNA replication.

Having observed a ~2-fold decrease in VACV DNA accumulation, but a 1.3 log decrease in late gene expression, we reasoned that the step in between the two may be the target of acrylamide-mediated inhibition. To quantify the effect of acrylamide on VACV mRNA levels we performed qRT-PCR on canonical early (J2R), intermediate (G8R) and late (F17) genes (Huttunen and Mercer 2019). Again, AraC-treated infected cells were used to benchmark inhibition of intermediate and late gene transcription. In both AraC and acrylamide treated conditions we observed no significant change in early gene transcription relative to the DMSO-treated control, and as anticipated AraC abolished intermediate and late gene expression (Fig. 2G). In the presence of acrylamide, we observed a 3-fold and >20-fold drop in intermediate and late gene transcription, respectively (Fig. 2G). These results indicated acrylamide inhibited post-replication VACV gene transcription.

### Vimentin is not required during VACV infection

Risco *et al.* proposed that vimentin acts to coordinate replication site formation and virus assembly (Risco et al. 2002). As we saw a minor defect in replication site size and a primary defect in post-replication gene transcription using acrylamide, we sought to determine if vimentin played an additional role in virus assembly. To this end we generated a vimentin null HeLa cell line using CRISPR. The lack of vimentin expression was confirmed at the transcriptional level by qRT-PCR (Fig. 3A; top) and at the protein level by immunoblot (Fig. 3B; bottom). Comparison of the actin, tubulin and lamin networks in parental and vimentin-null cells, by immunofluorescence, revealed no major differences in network architecture, cell size or cell shape (Fig. 3C).

**Figure 3.**
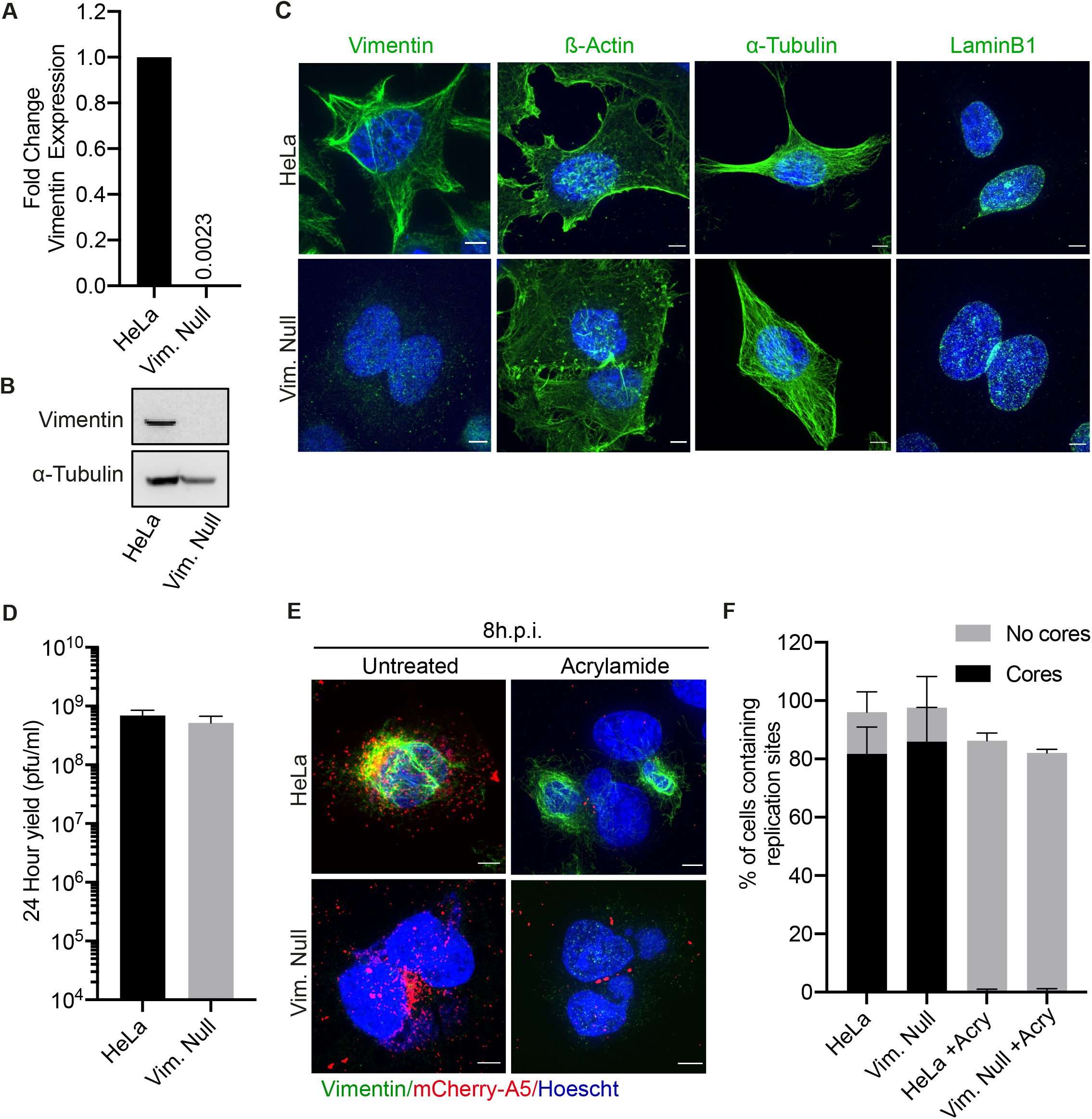
Acrylamide mediated inhibition of VACV infection is not due to collapsing of vimentin filaments. (A) Quantification of vimentin mRNA levels in parental HeLa and HeLa Vimentin Null (Vim. Null) cells using RTqPCR. GAPDH was used to normalise expression across all samples and the fold change vimentin mRNA calculated using threshold cycles. (B) Representative immunoblot of HeLa and Vim. Null cell lysates immunoblotted for vimentin; α-tubulin served as a loading control. (C) HeLa and Vim. Null cells were stained for cytoskeletal elements: α-Tubulin, LaminB1 or β-actin and DNA visualised using Hoechst. Scale bars = 5μm. (D) HeLa and Vim. Null cells were infected with WT WR at MOI 1 and viral yield determined at 24 hpi on BSC40 cells. (E) HeLa and Vim. Null cells were infected with WR mCh-A5 (red) at MOI 10 in the absence or presence of acrylamide. At 8 hpi cells were stained for vimentin (green), DNA (blue). Representative cell images shown. Scale bars = 5μm. (F) The percentage of cells containing replication sites, with and without mCh-A5 cores from (E) (n>75). All experiments were performed in triplicate with graphs representing the mean +1 SD.

Next we sought to determine the ability of the vimentin-null cells to support productive VACV infection using a 24 h yield assay. To our surprise we saw no difference VACV infectious yield between parental and vimentin-null cell lines (Fig. 3D). This result indicated that vimentin is not required for productive VACV infection. In light of this observation, we speculated that acrylamide-mediated inhibition of VACV might be due to the collapse of vimentin filaments constricting genome replication or impeding the spatial organisation of replication sites. To investigate this possibility, parental HeLa and vimentin-null cells were infected with WR mCh-A5 in the presence or absence of acrylamide. At 8hpi cells were fixed, nuclei and replication sites stained with Hoechst and immunostaining performed to visualise vimentin. In untreated HeLa cells vimentin was associated with replication sites and newly assembled virions were found throughout the cytoplasm (Fig. 3E). As expected, in acrylamide-treated HeLa cells vimentin was collapsed around replication sites and no virions were formed. Consistent with the 24h yield results, both replication sites and newly formed virions were seen in vimentin-null cells (Fig. 3E). Contrary to our hypothesis however, treatment of vimentin-null cells with acrylamide blocked the production of virions as effectively as in parental HeLa cells. Quantification of this phenotype indicated that regardless of the presence of vimentin, acrylamide treatment causes a small reduction in the number of cells with replication sites while completely abolishing the formation of nascent VACV virion (Fig. 3F). Thus, using a vimentin-null cell line we demonstrate that inhibition of VACV infection by acrylamide is independent of its ability to collapse cell’s vimentin network.

### Acrylamide blocks translation and stimulates the formation of antiviral granules

In addition to collapsing vimentin, acrylamide has been reported to activate cellular oxidative and ER stress responses (Komoike and Matsuoka 2016, Komoike and Matsuoka 2019), which converge upon eIF2α phosphorylation and global translational arrest (Jackson et al. 2010, McCormick and Khaperskyy 2017). Translational arrest, in turn leads to the formation of cytosolic stress granules (SGs) (Guzikowski, Chen and Zid 2019). Similar to SGs, antiviral granules (AVGs), have been reported to form and restrict VACV gene expression. AVGs were reported to form in ~10% of WT infected cells, in cells infected with VACV strains that lack the ability to block PKR-mediated responses and upon the addition of small compounds or antivirals that increase cellular stress (Simpson-Holley et al. 2011, Rozelle et al. 2014).

To determine if acrylamide was stimulating the formation of AVGs, HeLa cells were infected with WR mCh-A5 in the absence or presence of acrylamide. At 8 hpi, cells were fixed, stained for nuclei and replication sites using Hoechst, and immunofluorescence performed for G3BP1 and eIF4G, two components of AVGs. In untreated cells, G3BP1 and eIF4G staining was diffuse throughout cells and largely excluded from viral replication sites (Fig. 4A). In acrylamide-treated infected cells both G3BP1 and eIF4G showed distinctive localisation to replication sites, consistent with the formation of stress-induced AVGs (Rozelle et al. 2014).

**Figure 4.**
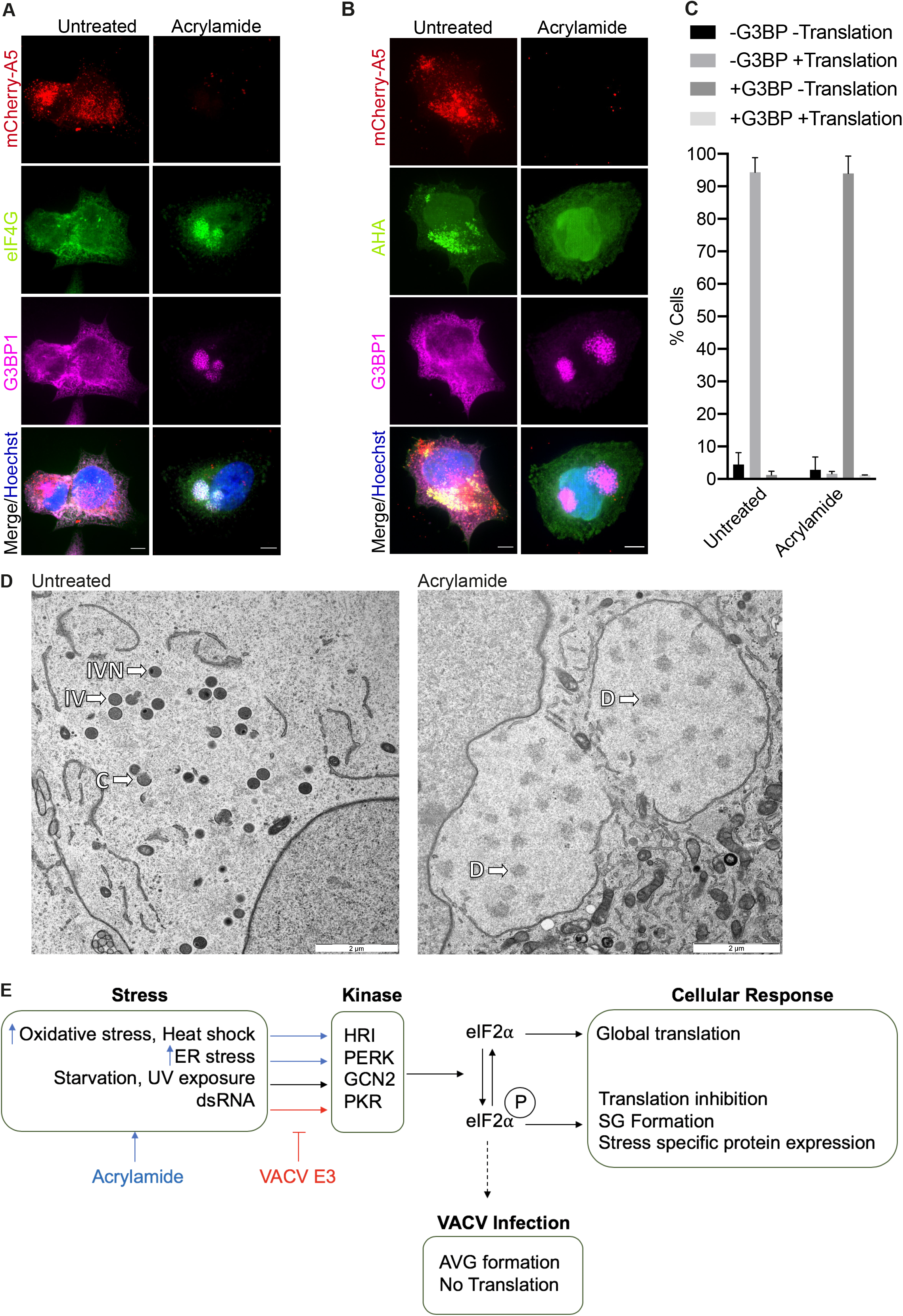
Acrylamide blocks translation and stimulates formation of anti-viral granules. (A) HeLa cells were infected with WR mCh-A5 (red) at MOI 10 in the absence or presence of acrylamide. At 8 hpi cells were fixed and stained for eIF4G (green) and G3BP1 (magenta), or DNA (blue). Scale bars = 5μm. (B) HeLa cells were infected as in (A) and AHA added 30 minutes prior to fixation at 8 hpi. AHA metabolic labelling was visualised using click-it reaction (green) followed by staining for G3BP (magenta) and DNA (blue). Scale bars = 5μm. (C) Cells from (B) were scored and quantified for the presence or absence of G3BP1 and active translation (AHA). Graph represent the mean + 1SD (n>75). (D) HeLa cells were infected with WT WR at MOI 10 in the absence or presence of acrylamide. At 8 hpi cells were fixed and processed for electron microscopy. Representative images of replication sites containing viral intermediates. (C = crescent, IV = intermediate virion, IVN = intermediate virion with nucleoid, D = dense region). All experiments were performed in triplicate.

As AVGs are known to form when VACV translation is blocked, we next looked at nascent protein synthesis. Again, cells were infected with WR mCh-A5 in the absence or presence of acrylamide. To visualize active translation the amino acid analogue Click-iT l-azidohomoalanine (AHA) was added to cells 30 min prior to fixation at 8 hpi. Cells were stained for nuclei and replication sites using Hoechst, and immunofluorescence performed for G3BP1. In untreated cells, active translation (AHA) was clearly seen within VACV replication sites while G3BP1 remained diffusely spread throughout the cell (Fig. 4B). The converse was seen in the acrylamide treated cells, in which negligible levels of active translation were observed and G3BP1 strongly localised to VACV replication sites.

To quantify this phenotype, each cell was scored as containing clear G3BP1 puncta within replication sites or not (+G3BP/−G3BP) (Fig. 4C). These two categories were then divided into cells with evident AHA accumulation within the replication site and those without (+translation/-translation). When the percentage of cells within these 4 subcategories was determined, two clear cell populations emerged: in the untreated cells the majority of replication sites were translation (AHA) positive and G3BP1 (AVG) negative, whereas in acrylamide-treated cells most replication sites were translation (AHA) negative and G3BP1 (AVG) positive (Fig. 4C). These results are consistent with previous reports that G3BP1 localisation to replication sites correlates with a lack of viral gene translation (Rozelle et al. 2014).

### Acrylamide treatment blocks infection prior to the initiation of VACV morphogenesis

As the acrylamide-mediated block of VACV infection was not related to vimentin-assisted formation of replication sites and virions, we wanted to visualize to impact of acrylamide on these viral structures. HeLa cells were infected with WT VACV in the absence or presence of acrylamide and processed for transmission electron microscopy at 8 hpi. In the untreated control sample, viral intermediates including crescents (C), immature virions (IVs) and immature virions with nucleoid (IVN) were observed (Fig. 4D). In the presence of acrylamide however, early viral replication sites wrapped in endoplasmic reticulum (Punjabi and Traktman 2005) were seen. While with no discernible viral intermediates were present, the replication sites contained areas of increased density (D). These results confirm and extend our finding that acrylamide inhibits the assembly of nascent VACV virions through inhibition of late protein synthesis, a prerequisite of virion assembly.

## Discussion

The complex cytoplasmic lifecycle of VACV has been the subject of intense study for nearly 70 years (Moss 2013). Despite this, the contribution of cellular factors to morphogenesis of poxvirus particles is scarcely defined (Condit, Moussatche and Traktman 2006). The intermediate filament vimentin was an exception, having been found in viral replication sites and virions, it was suggested to coordinate the assembly of both (Risco et al. 2002, Resch et al. 2007), (Manes et al. 2008). As vimentin is important for cytosolic organisation, maintenance of the microtuble network and cell integrity (Lowery et al. 2015, Ivaska et al. 2007), we wanted to better undertsand its role in VACV replication and assembly.

Following on from studies investigating the role of vimentin in virus infection, we employed acrylamide to collapse the vimentin filament network (Bhattacharya et al. 2007, Cordo and Candurra 2003). While known to have multiple effects, in the absence of commercially available vimentin polymerisation inhibitors, acrylamide is a useful tool for studying the role of these intermediate filaments (Durham et al. 1983). We established that treatment of infected cells with acrylamide reduced VACV yields by >99%. Refining the stage of virus infection, we found partial inhibition of intermediate gene expression and genome replication culminating in a total block late gene expression and subsequent virus assembly.

To verify that acrylamide mediated inhibition of VACV replication was due to disruption of the vimentin network we generated a vimentin-null HeLa cell line. We found that the absence of vimentin did not impact VACVs ability to form replication sites or to produce new infectious particles. We explored the possibility that the collapse of vimentin filaments around VACV replication sites, rather than its absence or general disruption, could be responsible for the block in virus assembly seen in the presence of acrylamide. However, treatment of VACV infected vimentin-null cells with acrylamide still blocked infection. This finding diverges from other studies that used acrylamide to dissect the role of vimentin in virus infection (Issac et al. 2014, Miller and Hertel 2009, Fay and Pante 2013, Cordo and Candurra 2003, Bhattacharya et al. 2007).

Despite our finding that vimentin removal has no impact upon VACV replication, it is found in replication sites and is packaged into VACV virions. Consistent with the role of this intermediate filament in cellular organelle positioning (Minin and Moldaver 2008) perhaps vimentin plays a supportive, as opposed to essential, role in the VACV lifecycle. It will be of future interest to explore the organisation of viral replication sites or virion sub-structures, in particular LBs, in the absence of vimentin.

Having uncoupled the effect of acrylamide on VACV infection from its ability to collapse the vimentin network, we turned our attention to other effects of acrylamide. A documented outcome of acrylamide exposure is the induction of oxidative and ER stress response pathways leading to the phosphorylation of eIF2α (Komoike and Matsuoka 2016, Komoike and Matsuoka 2019). The consequence of this is translational arrest, which can lead to the formation of cytosolic membraneless organelles termed stress granules (SGs) (Guzikowski et al. 2019). SGs have been show to play a role in cellular-antiviral response by inhibiting the accumulation of viral proteins upon oxidative stress (signalling via HRI kinase), ER stress (signalling via PERK), and upon sensing of dsRNA (signalling via PKR) (McEwen et al. 2005, Galabru and Hovanessian 1987, Harding et al. 2000, Piotrowska et al. 2010, Farrell et al. 1977, Onomoto et al. 2014). In turn, viruses have evolved mechanisms to manipulate stress granule formation and associated signalling pathways (McCormick and Khaperskyy 2017).

Antiviral granules (AVGs) are inhibitory cytoplasmic formations which share similarities to SGs and form spontaneously at low levels during normal VACV infection, or at higher levels when cells are subject to stimuli such as oxidative stress or altered RNA helicase activity (Simpson-Holley et al. 2011, Rozelle et al. 2014, Liem and Liu 2016). To counteract AVG formation VACV, encodes the E3 protein that binds to and masks dsRNA thereby prevent host recognition and PKR activation (Watson, Chang and Jacobs 1991, Chang, Watson and Jacobs 1992). Given the block in post-replicative gene expression, we investigated the possibility that acrylamide was inducing the formation of AVGs. Consistent with their formation, G3BP1 and eIF4G localised to nearly 100% of replication sites in acrylamide-treatment infected cells. Furthermore, active translation was negligible within these AVG containing replication sites.

As illustrated in Figure 4E, we propose that acrylamide activates non-PKR mediated cellular stress response pathway(s) which promote eIF2 phosphorylation and the formation of AVGs. This results in the slowing of viral genome replication and post-replicative intermediate transcription. VACV gene expression occurs in temporal waves with intermediate genes serving as late transcription factors. Thus the inhibition of viral translation by AVGs has a knock-on effect leading to reduced late gene transcription/translation and culminating in the lack of late gene products required for assembly of nascent VACV virions. These results suggest, consistent with previous reports (Rozelle et al. 2014), that exogenous stimulation of AVGs formation may be a useful strategy to promote cellular antiviral responses that bypass mechanisms used by VACV to manipulate host cell signalling.

## Materials and Methods

### Cell culture

BSC40 (African green monkey), HeLa and vimentin null HeLa cells were maintained at 37°C and 5% CO2 in complete Dulbecco’s modified Eagle’s medium (DMEM) containing 10% foetal bovine serum (FBS), 2mM GlutaMAX and 1% penicillin-streptomycin. The vimentin null cell line was generated by transfection of a vimentin Double Nickase plasmid mix (Santa- Cruz sc-400035-NIC) using Lipofectamine 2000 as per the manufacturer’s instruction. Transfected cells were initially selected using puromycin (1μg/ml) before single colonies were selected and assayed.

### Antibodies and reagents

The following antibodies were used for immunofluorescence; anti-vimentin (Abcam ab20346, 1:250), anti-vimentin (Sigma-Aldrich V5255, 1:200), anti-eIF4G1 (abcam ab47649,1:400), anti-G3BP (abcam ab56574, 1:400), anti-α-tubulin (Cell signalling technology 3873S, 1:1000), anti-LaminB1 (Abcam ab133741, 1:200), anti-β-Actin (Sigma-Aldrich A1978, 1:400), Alexa Flour secondary antibodies (Invitrogen, 1:500). For western blotting the following antibodies were used; anti-A10 (Immune Technology IT-012-010M1, 1:2000), anti-A17 (1:1000), anti-F17 (1:2500), anti-vimentin (Sigma-Aldrich V5255, 1:1000), anti-α-tubulin (Cell signalling technology 3873S, 1:5000). Drugs were used at the following concentrations; 10μM Cytosine arabinoside (AraC), 50μM MG132, 3μg/ml actinomycin D (ActD), 1mM cycloheximide (CHX), 30μM nocodazole, 4mM acrylamide.

### Viruses, VACV purification and infections

All viruses used in this study were derived from the VACV western reserve (WR) strain. In addition to wild type (WT WR), recombinant WR mCherry-A5 (Schmidt et al. 2011), WR early EGFP (WR E-GFP) and WR late EGFP (WR L-GFP) viruses (Kilcher et al. 2014) were used as described previously. Viruses were purified by sedimentation through a sucrose cushion. Briefly, infected BSC40 cells were scraped into PBS and harvested at 300g for 5 mins. The pellet was then resuspended in 10 mM Tris, pH 9.0 and incubated on ice for 5 mins. Cells were disrupted in a tight fitting Dounce homogeniser before centrifugation at 2000g, 10 min, 4°C. The clarified supernatant was collected, and this centrifugation step repeated. The supernatant was then loaded over 36% sucrose in 20 mM Tris, pH 9.0 and then harvested at 38,000g 4°C for 80 mins. The resulting pellet was resuspended in 200μl 1 mM Tris and stored at −80°C, titres were determined by plaque assay as described below.

Infections were carried out at the specified MOIs in minimal volumes of serum free DMEM. VACV was thawed and virions resuspended with several rounds of sonication in a water bath. Virus dilutions were incubated on cell monolayers for 1 hour with agitation every 15 minutes. Cells were then fed with complete DMEM and incubated in conditions as indicated.

### 24-hour yield and plaque assays

Confluent cells were infected with the appropriate viruses in 60mm dishes at an MOI of 1 for 24 hours. Cells were then scraped into PBS and harvested at 700g, 4°C for 5min. The cell pellet was suspended in 100μl 1mM Tris pH 9.0 and subjected to three rounds of freeze thawing in liquid nitrogen. Plaque assays were performed by titrating harvested virus onto confluent monolayers of BSC40 cells in 6 well dishes. Cells were infected with ten-fold dilutions of VACV and incubated for 48 hours before being stained with 0.1% crystal violet in 3.7% PFA.

### Immunofluorescence

For immunofluorescence analysis, DMEM was removed and cells were fixed in 4% (v/v) paraformaldehyde in PBS and permeabilised in 0.2% (v/v) Triton X-100 in PBS for 15 minutes each, with three PBS washes after each incubation. Alternatively, cells were fixed and permeabilised by incubating on ice with cold methanol for 10 minutes. Bovine serum albumin (BSA) at 1% (w/v) in PBS was used as a blocking agent and incubated with the cells for a minimum of 1 hour. Coverslips were incubated with primary antibodies diluted in blocking agent for 1 hour at indicated concentrations. Secondary antibodies were diluted in blocking agent and incubated with the cells for 1 hour. Coverslips were stained with Hoechst (1μg/ml in PBS) for 15mins before mounting with Immu-Mount (Thermo Scientific) and sealing with nail polish. Five washing steps were performed after each staining step and all incubation steps were carried out at room temperature, protected from light unless otherwise stated. Samples were visualised using Nikon eclipse Ti iSIM with NIS-Elements AR software and Images analysed using Image J 2.0.

### Visualisation of Nascent Protein synthesis

Metabolic labelling and visualisation were done using Click-iT^®^ AHA (L-azidohomoalaine) and the Click-iT™ Cell Reaction Buffer Kit (Invitrogen), according to the manufacturer’s instructions. Briefly, cells to be analysed were incubated with DMEM containing no glutamine, methionine or cystine for 1 hour prior to fixation. AHA was then added to cells at a concentration of 50μM for an additional 30 mins before fixation with 4% PFA and permeabilisation with 0.2% Triton for 15 minutes each. The click- it reaction cocktail was prepared containing an Alexa Fluor 488 Alkyne and incubated on coverslips for 30mins protected from the light. Coverslips were then washed and processed as per the immunofluorescence protocol above.

### Flow cytometry analysis of VACV gene expression

HeLa cells were seeded into 24 well dishes, treated with appropriate drugs and infected at MOI 4 with either WR E-GFP, or WR L-GFP viruses for 6 and 8 hours respectively. For analysis by flow cytometry, cells were washed with PBS and removed from the plate surface by trypsin treatment. Equal volumes of blocking buffer (5% FBS in PBS) and PFA (4% final concentration in PBS) were added to each well and incubated for 15 minutes. Samples were then transferred to 96 well plates ready for analysis by flow cytometry. All solutions were prewarmed to 37°C and all conditions were performed in triplicate. The Guava easyCyte HT instrument was used in conjunction with the Guava soft Incyte 3.1 software for analysis.

### RT-qPCR

For analysis of viral gene expression, HeLa cells were grown in 6 well dishes and infected in triplicate with WT WR at a MOI of 10 for either 2 hours (early), 4 hours (intermediate) or 8 hours (late). For CRISPR analysis, HeLa and vimentin null cells were grown to confluency in 6 well dishes before harvesting. The Qiagen RNeasy kit was used to extract RNA according to the manufacturer’s instructions. RNA concentration was measured using the Nanodrop 2000 Spectrophotometer and 1μg per sample was used as template for reverse transcription. cDNA was generated using SuperScript II reverse transcriptase (Thermo Fisher Scientific) and oligo(dT) primers. qPCR reactions were carried out using Mesa Blue qPCR MasterMix and appropriate primers on the BioRad CFX connect qPCR machine. Early samples were analysed with VACV J2R (5⍰-TACGGAACGGGACTATGGAC-3⍰ and 5⍰-GTTTGCCATACGCTCACAGA-3⍰) primers, intermediate with G8R (5⍰-AATGTAGACTCGACGGATGAGTTA-3⍰ and 5⍰-TCGTCATTATCCATTACGATTCTAGTT-3⍰) primers, late with F17R (5⍰-ATTCTCATTTTGCATCTGCTC-3⍰ and 5⍰-AGCTACATTATCGCGATTAGC-3⍰) primers and vimentin with (5⍰-GCTCGTCACCTTCGTGAATA-3⍰ and 5⍰-CAGAGGGAGTGAATCCAGATTAG-3⍰) primers. GAPDH primers were used on all samples as a housekeeping control. Fold change in mRNA levels was calculated using threshold cycle (C_T_) values normalised against the housekeeping control.

### Quantification of viral DNA by qPCR

VACV genome replication was quantified by qPCR as previously described (Huttunen and Mercer 2019). In short, cells were scraped into PBS and harvested at 400g for 5 mins. Genomic DNA was then extracted using the DNeasy Blood & Tissue Kit (Qiagen) as per manufacturer’s instructions. A VACV genomic DNA dilution series of known concentration was used to create a standard curve. Samples were analysed by qPCR using the Mesa Blue qPCR MasterMix and C11R primer (5⍰-AAACACACACTGAGAAACAGCATAAA-3⍰, 5⍰-ACTATCGGCGAATGATCTGATTA-3⍰) BioRad CFX connect qPCR machine.

### Electron Microscopy

HeLa cells on coverslips were infected at MOI 10 with WT WR and treated as indicated. At 8hpi the coverslips were fixed in EM-grade 2% paraformaldehyde/2% glutaraldehyde (TAAB Laboratories Equipment, Ltd.) in 0.1 M sodium cacodylate, secondarily fixed for 1 h in 1% osmium tetraoxide/1.5% potassium ferricyanide at 4°C and then treated with 1% tannic acid in 0.1M sodium cacodylate for 45min at room temperature. Samples were then dehydrated in sequentially increasing concentration of ethanol solutions, and embedded in Epon resin. Coverslips were inverted onto prepolymerised Epon stubs and polymerised by baking at 60°C overnight. The 70nm thin sections were cut with a Diatome 45° diamond knife using an ultramicrotome (UC7; Leica). Sections were collected on 1 ×2mm formvar-coated slot grids and stained with Reynolds lead citrate. All samples were imaged using a transmission electron microscope (Tecnai T12; FEI) equipped with a charge-coupled device camera (SIS Morada; Olympus).

### Virus fractionations and western blotting

Purified WT WR virus was pelleted at 16,000 g, room temperature for 30 min and resuspended in fractionation buffer (1% NP-40, 50 mM DTT in 1 mM Tris pH 9.0). After incubation at 37°C for 30 min, samples were centrifuged at 16,000 g, 4°C for 30 min. Pellets (core and lateral body fraction) were resuspended in 10mM Tris pH 9.0 and supernatants (membrane fraction) transferred to a fresh tube. To remove lateral bodies from cores, the combined fraction was resuspended in varying concentrations of trypsin (from 0.125 to 1 μg/ml made up in 10mM Tris pH 9.0) and incubated at 37°C for 15 minutes. Trypsin inhibitor was then added followed by a 15 min incubation at 25°C. Samples were centrifuged at 15,000 rpm, 4°C for 30 min, the supernatant was transferred to a fresh tube and the core containing pellets resuspended in 10mM Tris pH 9.0. For analysis, samples were boiled with Laemmli buffer and separated on SDS-PAGE gels prior to transfer of proteins onto nitrocellulose membranes. Proteins were visualized using indicated antibodies on the Li-Cor Odyssey 3.0 imaging system.

## Acknowledgements

We thank Prof. Paula Traktman (Medical University of South Carolina) for A17 and F17 antisera. I.J. White is supported by MRC core funding to the MRC Laboratory for Molecular Cell Biology at University College London, award code (MC_U12266B).

## Contributions

J.J. Wood and J. Mercer designed the study. J.J. Wood performed and analysed experiments. I.J. White prepared and imaged EM samples. J.J. Wood and J. Mercer wrote the manuscript with input from I.J. White.

## References

Balzarini, J. & E. Declercq (1989) THE ANTIVIRAL ACTIVITY OF 9-BETA-D-ARABINOFURANOSYLADENINE IS ENHANCED BY THE 2’,3’-DIDEOXYRIBOSIDE, THE 2’,3’-DIDEHYDRO-2’,3’-DIDEOXYRIBOSIDE AND THE 3’-AZIDO-2’,3’-DIDEOXYRIBOSIDE OF 2,6-DIAMINOPURINE. Biochemical and Biophysical Research Communications, 159, 61–67.

Bhattacharya, B., R. J. Noad & P. Roy (2007) Interaction between Bluetongue virus outer capsid protein VP2 and vimentin is necessary for virus egress. Virology Journal, 4.

Bidgood, S. R. (2019) Continued poxvirus research: From foe to friend. Plos Biology, 17.

Chang, H. W., J. C. Watson & B. L. Jacobs (1992) THE E3L GENE OF VACCINIA VIRUS ENCODES AN INHIBITOR OF THE INTERFERON-INDUCED, DOUBLE-STRANDED RNA-DEPENDENT PROTEIN-KINASE. Proceedings of the National Academy of Sciences of the United States of America, 89, 4825–4829.

Chang, L. & R. D. Goldman (2004) Intermediate filaments mediate cytoskeletal crosstalk. Nature Reviews Molecular Cell Biology, 5, 601–613.

Chung, C. S., C. H. Chen, M. Y. Ho, C. Y. Huang, C. L. Liao & W. Chang (2006) Vaccinia virus proteome: Identification of proteins in vaccinia virus intracellular mature virion particles. Journal of Virology, 80, 2127–2140.

Condit, R. C., N. Moussatche & P. Traktman (2006) In a nutshell: Structure and assembly of the vaccinia virion. Advances in Virus Research, Vol 66, 66, 31–+.

Cordo, S. M. & N. A. Candurra (2003) Intermediate filament integrity is required for Junin virus replication. Virus Research, 97, 47–55.

Cudmore, S., P. Cossart, G. Griffiths & M. Way (1995) ACTIN-BASED MOTILITY OF VACCINIA VIRUS. Nature, 378, 636–638.

Danielsson, F., M. K. Peterson, H. C. Araujo, F. Lautenschlager & A. K. B. Gad (2018) Vimentin Diversity in Health and Disease. Cells, 7.

Durham, H. D., S. D. J. Pena & S. Carpenter (1983) THE NEUROTOXINS 2,5-HEXANEDIONE AND ACRYLAMIDE PROMOTE AGGREGATION OF INTERMEDIATE FILAMENTS IN CULTURED FIBROBLASTS. Muscle & Nerve, 6, 631–637.

Eckert, B. S. (1986) ALTERATION OF THE DISTRIBUTION OF INTERMEDIATE FILAMENTS IN PTK1 CELLS BY ACRYLAMIDE.2. EFFECT ON THE ORGANIZATION OF CYTOPLASMIC ORGANELLES. Cell Motility and the Cytoskeleton, 6, 15–24.

Farrell, P. J., K. Balkow, T. Hunt, R. J. Jackson & H. Trachsel (1977) PHOSPHORYLATION OF INITIATION-FACTOR ELF-2 AND CONTROL OF RETICULOCYTE PROTEIN-SYNTHESIS. Cell, 11, 187–200.

Fay, N. & N. Pante (2013) The intermediate filament network protein, vimentin, is required for parvoviral infection. Virology, 444, 181–190.

Galabru, J. & A. Hovanessian (1987) AUTOPHOSPHORYLATION OF THE PROTEIN-KINASE DEPENDENT ON DOUBLE-STRANDED-RNA. Journal of Biological Chemistry, 262, 15538–15544.

Guzikowski, A. R., Y. S. Chen & B. M. Zid (2019) Stress-induced mRNP granules: Form and function of processing bodies and stress granules. Wiley Interdisciplinary Reviews-Rna, 10.

Harding, H. P., Y. H. Zhang, A. Bertolotti, H. Q. Zeng & D. Ron (2000) Perk is essential for translational regulation and cell survival during the unfolded protein response. Molecular Cell, 5, 897–904.

Huttunen, M. & J. Mercer. 2019. Quantitative PCR-Based Assessment of Vaccinia Virus RNA and DNA in Infected Cells. Vaccinia Virus. Methods in Molecular Biology: Humana, New York, NY.

Ichihashi, Y., M. Oie & T. Tsuruhara (1984) LOCATION OF DNA-BINDING PROTEINS AND DISULFIDE-LINKED PROTEINS IN VACCINIA VIRUS STRUCTURAL ELEMENTS. Journal of Virology, 50, 929–938.

Issac, T. H. K., E. L. Tan & J. J. H. Chu (2014) Proteomic profiling of chikungunya virus-infected human muscle cells: Reveal the role of cytoskeleton network in CHIKV replication. Journal of Proteomics, 108, 445–464.

Ivaska, J., H. M. Pallari, J. Nevo & J. E. Eriksson (2007) Novel functions of vimentin in cell adhesion, migration, and signaling. Experimental Cell Research, 313, 2050–2062.

Jackson, R. J., C. U. T. Hellen & T. V. Pestova (2010) The mechanism of eukaryotic translation initiation and principles of its regulation. Nature Reviews Molecular Cell Biology, 11, 113–127.

Jiang, L. P., J. Cao, Y. An, C. Y. Geng, S. X. Qu, L. J. Jiang & L. F. Zhong (2007) Genotoxicity of acrylamide in human hepatoma G2 (HepG2) cells. Toxicology in Vitro, 21, 1486–1492.

Kilcher, S., F. I. Schmidt, C. Schneider, M. Kopf, A. Helenius & J. Mercer (2014) siRNA Screen of Early Poxvirus Genes Identifies the AAA+ ATPase D5 as the Virus Genome-Uncoating Factor. Cell Host & Microbe, 15, 103–112.

Kim, K. H., B. Park, D. K. Rhee & S. Pyo (2015) Acrylamide Induces Senescence in Macrophages through a Process Involving ATF3, ROS, p38/JNK, and a Telomerase-Independent Pathway. Chemical Research in Toxicology, 28, 71–86.

Komoike, Y. & M. Matsuoka (2016) Endoplasmic reticulum stress-mediated neuronal apoptosis by acrylamide exposure. Toxicology and Applied Pharmacology, 310, 68–77.

Komoike, Y. & M. Matsuoka (2019) In vitro and in vivo studies of oxidative stress responses against acrylamide toxicity in zebrafish. Journal of Hazardous Materials, 365, 430–439.

Liem, J. & J. Liu (2016) Stress Beyond Translation: Poxviruses and More. Viruses-Basel, 8.

Lowery, J., E. R. Kuczmarski, H. Herrmann & R. D. Goldman (2015) Intermediate Filaments Play a Pivotal Role in Regulating Cell Architecture and Function. Journal of Biological Chemistry, 290, 17145–17153.

Mallardo, M., S. Schleich & J. K. Locker (2001) Microtubule-dependent organization of vaccinia virus core - derived early mRNAs into distinct cytoplasmic structures. Molecular Biology of the Cell, 12, 3875–3891.

Manes, N. P., R. D. Estep, H. M. Mottaz, R. J. Moore, T. R. W. Clauss, M. E. Monroe, X. Du, J. N. Adkins, S. W. Wong & R. D. Smith (2008) Comparative proteomics of human monkeypox and vaccinia intracellular mature and extracellular enveloped virions. Journal of Proteome Research, 7, 960–968.

McCormick, C. & D. A. Khaperskyy (2017) Translation inhibition and stress granules in the antiviral immune response. Nature Reviews Immunology, 17, 647–660.

McEwen, E., N. Kedersha, B. B. Song, D. Scheuner, N. Gilks, A. P. Han, J. J. Chen, P. Anderson & R. J. Kaufman (2005) Heme-regulated inhibitor kinase-mediated phosphorylation of eukaryotic translation initiation factor 2 inhibits translation, induces stress granule formation, and mediates survival upon arsenite exposure. Journal of Biological Chemistry, 280, 16925–16933.

Mercer, J. & A. Helenius (2008) Vaccinia virus uses macropinocytosis and apoptotic mimicry to enter host cells. Science, 320, 531–535.

Mercer, J., S. Knebel, F. I. Schmidt, J. Crouse, C. Burkard & A. Helenius (2010) Vaccinia virus strains use distinct forms of macropinocytosis for host-cell entry. Proceedings of the National Academy of Sciences of the United States of America, 107, 9346–9351.

Mercer, J., B. Snijder, R. Sacher, C. Burkard, C. K. E. Bleck, H. Stahlberg, L. Pelkmans & A. Helenius (2012) RNAi Screening Reveals Proteasome- and Cullin3-Dependent Stages in Vaccinia Virus Infection. Cell Reports, 2, 1036–1047.

Mercer, J. & P. Traktman (2003) Investigation of structural and functional motifs within the vaccinia virus A14 phosphoprotein, an essential component of the virion membrane. Journal of Virology, 77, 8857–8871.

Miller, M. S. & L. Hertel (2009) Onset of Human Cytomegalovirus Replication in Fibroblasts Requires the Presence of an Intact Vimentin Cytoskeleton. Journal of Virology, 83, 7015–7028.

Minin, A. A. & M. V. Moldaver (2008) Intermediate Vimentin Filaments and Their Role in Intracellular Organelle Distribution. Biochemistry-Moscow, 73, 1453–1466.

Moss, B. 2013. Poxviridae. Fields Virology, 6th ed, vol 2. Lippincott Williams and Wilkins, Philadelphia, PA.

Onomoto, K., M. Yoneyama, G. Fung, H. Kato & T. Fujita (2014) Antiviral innate immunity and stress granule responses. Trends in Immunology, 35, 420–428.

Piotrowska, J., S. J. Hansen, N. Park, K. Jamka, P. Sarnow & K. E. Gustin (2010) Stable Formation of Compositionally Unique Stress Granules in Virus-Infected Cells. Journal of Virology, 84, 3654–3665.

Ploubidou, A., V. Moreau, K. Ashman, I. Reckmann, C. Gonzalez & M. Way (2000) Vaccinia virus infection disrupts microtubule organization and centrosome function. Embo Journal, 19, 3932–3944.

Punjabi, A. & P. Traktman (2005) Cell biological and functional characterization of the vaccinia virus F10 kinase: Implications for the mechanism of virion morphogenesis. Journal of Virology, 79, 2171–2190.

Resch, W., K. K. Hixson, R. J. Moore, M. S. Lipton & B. Moss (2007) Protein composition of the vaccinia virus mature virion. Virology, 358, 233–247.

Rietdorf, J., A. Ploubidou, I. Reckmann, A. Holmstrom, F. Frischknecht, M. Zettl, T. Zimmermann & M. Way (2001) Kinesin-dependent movement on microtubules precedes actin-based motility of vaccinia virus. Nature Cell Biology, 3, 992–1000.

Risco, C., J. R. Rodriguez, C. Lopez-Iglesias, J. L. Carrascosa, M. Esteban & D. Rodriguez (2002) Endoplasmic reticulum-Golgi intermediate compartment membranes and vimentin filaments participate in vaccinia virus assembly. J Virol, 76, 1839–55.

Rizopoulos, Z., G. Balistreri, S. Kilcher, C. K. Martin, M. Syedbasha, A. Helenius & J. Mercer (2015) Vaccinia Virus Infection Requires Maturation of Macropinosomes. Traffic, 16, 814–831.

Rozelle, D. K., C. M. Filone, N. Kedersha & J. H. Connor (2014) Activation of Stress Response Pathways Promotes Formation of Antiviral Granules and Restricts Virus Replication. Molecular and Cellular Biology, 34, 2003–2016.

Schmidt, F. I., C. K. E. Bleck, A. Helenius & J. Mercer (2011) Vaccinia extracellular virions enter cells by macropinocytosis and acid-activated membrane rupture. Embo Journal, 30, 3647–3661.

Schmidt, F. I., C. K. E. Bleck, L. Reh, K. Novy, B. Wollscheid, A. Helenius, H. Stahlberg & J. Mercer (2013) Vaccinia Virus Entry Is Followed by Core Activation and Proteasome-Mediated Release of the Immunomodulatory Effector VH1 from Lateral Bodies. Cell Reports, 4, 464–476.

Sidwell, R. W., G. J. Dixon, S. M. Sellers & F. M. Schabel (1968) IN VIVO ANTIVIRAL PROPERTIES OF BIOLOGICALLY ACTIVE COMPOUNDS.2. STUDIES WITH INFLUENZA AND VACCINIA VIRUSES. Applied Microbiology, 16, 370-&.

Simpson-Holley, M., N. Kedersha, K. Dower, K. H. Rubins, P. Anderson, L. E. Hensley & J. H. Connor (2011) Formation of Antiviral Cytoplasmic Granules during Orthopoxvirus Infection. Journal of Virology, 85, 1581–1593.

Smith, G. L. & M. Law (2004) The exit of Vaccinia virus from infected cells. Virus Research, 106, 189–197.

Styers, M. L., G. Salazar, R. Love, A. A. Peden, A. P. Kowalczyk & V. Faundez (2004) The endo-lysosomal sorting machinery interacts with the intermediate filament cytoskeleton. Molecular Biology of the Cell, 15, 5369–5382.

Wang, Q., X. L. Zhang, Y. L. Han, X. L. Wang & G. X. Gao (2016) M2BP inhibits HIV-1 virion production in a vimentin filaments-dependent manner. Scientific Reports, 6.

Ward, B. M. & B. Moss (2001) Vaccinia virus intracellular movement is associated with microtubules and independent of actin tails. Journal of Virology, 75, 11651–11663.

Watson, J. C., H. W. Chang & B. L. Jacobs (1991) CHARACTERIZATION OF A VACCINIA VIRUS-ENCODED DOUBLE-STRANDED RNA-BINDING PROTEIN THAT MAY BE INVOLVED IN INHIBITION OF THE DOUBLE-STRANDED RNA-DEPENDENT PROTEIN-KINASE. Virology, 185, 206–216.

Yakimovich, A., M. Huttunen, B. Zehnder, L. J. Coulter, V. Gould, C. Schneider, M. Kopf, C. J. McInnes, U. F. Greber & J. Mercer (2017) Inhibition of Poxvirus Gene Expression and Genome Replication by Bisbenzimide Derivatives. Journal of Virology, 91.

